# SpacePHARER: Sensitive identification of phages from CRISPR spacers in prokaryotic hosts

**DOI:** 10.1101/2020.05.15.090266

**Authors:** R. Zhang, M. Mirdita, E. Levy Karin, C. Norroy, C. Galiez, J. Söding

## Abstract

SpacePHARER (CRISPR Spacer Phage-Host Pair Finder) is a sensitive and fast tool for *de novo* prediction of phage-host relationships via identifying phage genomes that match CRISPR spacers in genomic or metagenomic data. SpacePHARER gains sensitivity by comparing spacers and phages at the protein level, optimizing its scores for matching very short sequences, and combining evidence from multiple matches, while controlling for false positives. We demonstrate SpacePHARER by searching a comprehensive spacer list against all complete phage genomes.

## I. INTRODUCTION

Viruses of bacteria and archaea (phages) are the most abundant biological entities in nature. However, little is known about their roles in the microbial ecosystem and how they interact with their hosts, as cultivating most phages and hosts in the lab is challenging. Many prokaryotes (40% of bacteria and 81% of archaea) possess an adaptive immune system against phages, the Clustered Regularly Interspaced Short Palindromic Repeat (CRISPR) system [6]. After surviving a phage infection, they can incorporate a short DNA fragment (28-42 nt) as a spacer in a CRISPR array. The transcribed spacer will be used with other Cas components for a targeted destruction of future invaders. Some CRISPR-Cas systems require a 2-6 nucleotide long, highly conserved protospacer-adjacent motif (PAM) flanking the viral target to prevent autoimmunity. Multiple spacers targeting the same invader are not uncommon, due to either multiple infection events or the primed spacer acquisition mechanism identified in some CRISPR subtypes. CRISPR spacers have been previously exploited to identify phage-host relationship [3, 11, 12, 15]. These methods compare individual CRISPR spacers with phage genomes using BLASTN [1] and apply stringent filtering criteria, e.g. allowing only up to two mismatches. They are thus limited to identifying very close matches. However, a higher sensitivity is crucial because phage reference databases are very incomplete and often will not contain phages highly similar to those to be identified. To increase sensitivity, (1) we compare protein coding sequences because phage genomes are mostly coding, and, to evade the CRISPR immune response, are under pressure to mutate their genome with minimal changes on the amino acid level; (2) we optimized a substitution matrix and gap penalties for short, highly similar protein fragments; (3) we combine evidence from multiple spacers matching to the same phage genome.

## II. METHODS

### Input

SpacePHARER accepts spacer sequences as multiple FASTA files each containing spacers from a single prokaryotic genome or as multiple output files from the CRISPR detection tools PILER-CR [7], CRT [5], MinCED [13] or CRISPRDetect [4]. Phage genomes are supplied as separate FASTA files or can be downloaded by SpacePHARER from NCBI GenBank [2]. Optionally, additional taxonomic labels can be provided for spacers or phages to be included in the final report.

### Algorithm

SpacePHARER is divided into five steps (**Figure 1A, Supp. Materials**). (0) Preprocess input: scan the phage genome and CRISPR spacers in six reading frames, extract and translate all putative coding fragments of at least 27 nt, with user-definable translation tables. Each query set *Q* consists of the translated ORFs *q* of CRISPR spacers extracted from one prokaryotic genome, and each target set *T* comprises the putative protein sequences *t* from a single phage. We refer to similar *q* and *t* as *hit*, and an identified host-phage relationship *Q* − *T* as *match*. (1) Search all *q*’s against all *t*’s using the fast, sensitive MMseqs2 protein search [14], with VTML40 substitution matrix [10], gap open cost of 16 and extension cost of 2 (Figure S1). We optimized a short, spaced k-mer pattern for the prefilter stage (10111011) with six informative (‘1’) positions. In addition, align all *q* − *t* hits reported in previous search on nucleotide level and prioritize near-perfect nucleotide hits (**Supp. Materials**). (2) For each *q* − *T* pair, compute the P-value for the best hit *p*_bh_ from first-order statistics. (3) Compute a combined score *S*_comb_ from best-hit P-values of multiple hits between *Q* and *T* using a modified truncated-product method (**Supp. Materials**). (4) Compute the false discovery rate (FDR = FP /(TP + FP)) and only retain matches with FDR *<* 0.05. For that purpose, SpacePHARER is run on a null model database and the fraction of null matches with *S*_comb_ below a cut-off (empirical P-value) is used to estimate the FDR. (5) Scan 10 nt upstream and downstream of the phage’s protospacer for a possible PAM.

**FIG 1.**
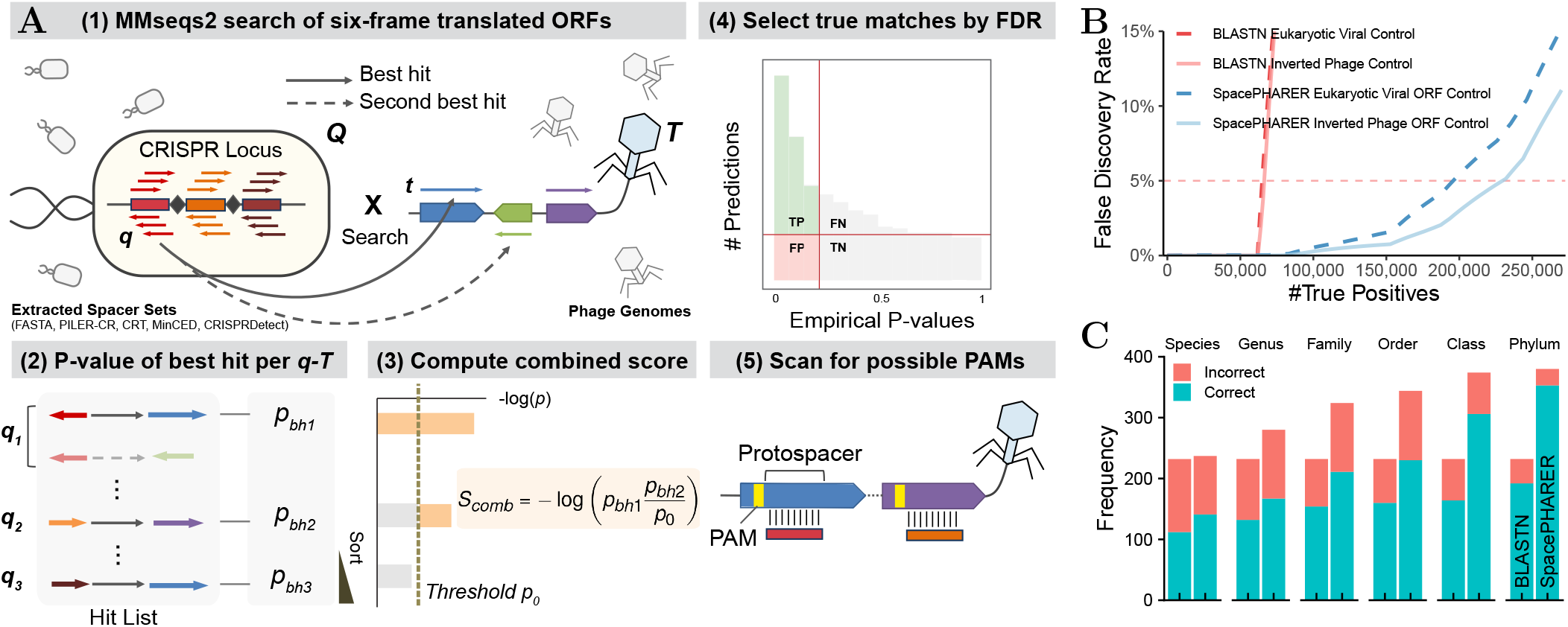
(**A**) SpacePHARER algorithm. A query set *Q* consists of 6-frame translated ORFs (*q*) from CRISPR spacers, and a target set *T* consists of 6-frame translated ORFs (*t*) of phage proteins. (1) Search all *q*s against all *t*s using MMseqs2. Align the *q* − *t* hits on nucleotide level and prioritize near-perfect nucleotide hits. (2) For each *q* − *T* pair, compute the P-value for the best hit from first-order statistics. (3) Compute score *S*_comb_ by combining the best-hit P-values from multiple hits between *Q* and *T* using a modified truncated-product method. (4) Estimate the FDR by searching a null database. (5) Scan for possible protospacer adjacent motif (PAM). (**B**) Performance comparison between SpacePHARER (blue) and BLASTN (red) using inverted phage sequences (solid lines) or eukaryotic viral ORFs as null set (dashed lines) demonstrated by expected number of true positive (TP) predictions at different false discovery rates (FDRs). (**C**) Performance comparison between BLASTN (left), SpacePHARER using the weighted lowest common ancestor procedure (LCA, right) at FDR = 0.02, evaluated by the number of correct (blue) and incorrect (red) predictions, for all the host predictions made at each taxonomic rank or below.

### Output

is a tab-separated text file. Each host-phage match spans two or more lines. The first starts with ‘#’: prokaryote accession, phage accession, *S*_comb_, number of hits in the match. Each following line describes an individual hit: spacer accession, phage accession, *p*_bh_, spacer start and end, phage start and end, possible 5’ PAM|3’ PAM, possible 5’ PAM|3’ PAM on the reverse strand. If requested, the spacer–phage alignments are included.

If taxonomic labels are provided, taxonomic reports based on the weighted lowest common ancestor (LCA) procedure described in [9] are created for host LCAs of each phage genome or phage LCAs of each spacer as additional tab-separated text files.

## III. RESULTS

### Datasets

We split a previously published spacer dataset [12] of 363,460 unique spacers from 30,389 prokaryotic genomes randomly into an optimization set (20%, 6,067 genomes) and a test set (80%, 24,322 genomes). The performance of SpacePHARER was evaluated on the spacer test set against a target database of 7,824 phage genomes. We used two null databases: 11,304 eukaryotic viral genomes and the inverted translated sequences of the target database. Viral genomes were downloaded from GenBank in 09/2018.

The performance of SpacePHARER in Figure 1C was evaluated on a validation dataset of spacers from 1,066 bacterial genomes against 809 phage genomes with annotated host taxonomy [8]. For each phage, we predicted the host based on the host LCA.

### Prediction quality

At FDR = 0.05, SpacePHARER predicted 3 to 4 times more prokaryote-phage matches than BLASTN (Figure 1B, Figure S2). SpacePHARER predicted the correct host for more phages than BLASTN at all taxonomic ranks, while including most of the BLASTN predictions, at better precision (Figure 1C, Figure S3). If the host or a close relative of a phage is absent in the database (either because the host is unidentified or the host lacks a CRISPR-Cas system), the predicted host may be correct only at a higher rank than species.

### Run time

SpacePHARER took 12 minutes to process the test dataset on 2 × 6-core 2.40 GHz CPUs, 47 times faster than BLASTN (575 minutes).

## IV. CONCLUSION

SpacePHARER is 1.4 to 4 × more sensitive than BLASTN in detecting phage-host pairs, due to searching with protein sequences, optimizing short sequence comparisons, and combining statistical evidence, and it is fast enough to analyze large-scale genomic and metagenomic datasets.

## Availability and implementation

SpacePHARER is available as an open-source (GPLv3), user-friendly command-line software for Linux and macOS: **spacepharer.soedinglab.org**.

## References

[1] Altschul, S.F. et al. (1990). Basic local alignment search tool. J. Mol. Biol., 215(3), 403–410.

[2] Benson, D.A. et al. (2013). GenBank. Nucleic Acids Res., 41(D1), D36–D42.

[3] Biswas, A. et al. (2013). CRISPRTarget: bioinformatic prediction and analysis of crRNA targets. RNA Biol., 10(5), 817–827.

[4] Biswas, A. et al. (2016). CRISPRdetect: A flexible algorithm to define CRISPR arrays. BMC Genom., 17(1), 356.

[5] Bland, C. et al. (2007). CRISPR recognition tool (CRT): a tool for automatic detection of clustered regularly interspaced palindromic repeats. BMC Bioinform., 8(1), 209.

[6] Burstein, D. et al. (2016). Major bacterial lineages are essentially devoid of crispr-cas viral defence systems. Nature Communications, 7(1), 10613.

[7] Edgar, R.C. (2007). PILER-CR: Fast and accurate identification of CRISPR repeats. BMC Bioinform., 8(1), 18.

[8] Edwards, R.A. et al. (2015). Computational approaches to predict bacteriophage–host relationships. FEMS Microbiol. Rev., 40(2), 258–272.

[9] Mirdita, M. et al. (2020). Fast and sensitive taxonomic assignment to metagenomic contigs. bioRxiv. doi:10.1101/2020.11.27.401018.

[10] Müller, T. et al. (2002). Estimating amino acid substitution models: A comparison of Dayhoff’s estimator, the resolvent approach and a maximum likelihood method. Mol. Biol. Evol., 19(1), 8–13.

[11] Paez-Espino, D. et al. (2016). Uncovering Earth’s virome. Nature, 536(7617), 425–430.

[12] Shmakov, S.A. et al. (2017). The CRISPR spacer space is dominated by sequences from species-specific mobilomes. mBio, 8(5), e01397–17.

[13] Skennerton, C. (2016). Minced - mining CRISPRs in environmental datasets. https://github.com/ctSkennerton/minced.

[14] Steinegger, M. and Söding, J. (2017). MMseqs2 enables sensitive protein sequence searching for the analysis of massive data sets. Nat. Biotechnol., 35(11), 1026–1028.

[15] Stern, A. et al. (2012). CRISPR targeting reveals a reservoir of common phages associated with the human gut microbiome. Genome Res., 22(10), 1985–1994.

