## Supplementary Material for "SpacePHARER: Sensitive identification of phages from CRISPR spacers in prokaryotic hosts"

### I. ALGORITHM DESCRIPTION

The query spacer set $Q$ has $N_q$ translated ORFs $q$ of CRISPR spacers ($Q = \{q_1, ...q_{N_q}\}$) from one prokaryotic genome. Phage proteome target set $T$ has $N_t$ phage protein sequences $t$ ($T = \{t_1, ...t_{N_t}\}$). These protein sequences are extracted in the input preprocessing step (Step 0) of the algorithm from each spacer set and each phage genome by scanning them in six translational frames. We refer to similarity between $q$ and $t$ as hit, and similarity between $Q$ and $T$ as match. The SpacePHARER algorithm relies on a statistic for the combination of hits between a spacer sequence set and a phage protein sequence set. The idea is that combining together several sub-significant hits (due to weak homologies or the typical length of spacers) can be highly informative and result in a significant match. Steps 2 and 3 of the algorithm test if the pairwise P-values of the best hit of sequences in the query set with those in the target set are due to homologous relationships or entirely due to chance.

#### A. (1) MMseqs2 protein-level search

The SpacePHARER algorithm first searches all $q$'s against all $t$'s using the fast, sensitive MMseqs2 protein-level search [10], with VTML40 substitution matrix [8], gap open cost of 16, gap extension cost of 2, and a short, spaced k-mer pattern for the prefilter stage (10111011) with six informative ("1") positions. Spaced k-mers are utilized in MMseqs2 to reduce the correlation between k-mers at neighboring positions, and to achieve better sensitivity and speed. The spaced k-mer pattern is chosen such that it is short in length in order to produce consecutive double k-mer matches (which are demanded by MMseqs2) within spacer fragments of 10-12 aa, and that the number of maximum overlapping informative positions is minimized.

Perfect or near-perfect hits (with no or 1-2 mismatches on the nucleotide level) are shown to be very reliable signals in predicting phage-host relationship and improve the taxonomic certainty of the prediction, even if there is only a single hit between a phage-host pair [4]. However, those hits are not well reflected in the pairwise P-value of the protein-level search. Therefore, all $q - t$ hits reported from the sensitive protein-level search will be aligned again on the nucleotide level with match reward of 1, mismatch penalty of 1, gap open cost of 10 and gap extension cost of 2. The protein-level search will compute a protein pairwise P-value ($p_{\text{prot}}$) for each hit and nucleotide alignment a nucleotide pairwise P-value ($p_{\text{nucl}}$). In order to prioritize near-perfect hits on the nucleotide level to gain precision without losing much sensitivity, we compute the pairwise P-value as

$$\exp\left(\min\left\{\left(0.5\log p_{prot} + 0.5\log p_{nucl}\right), \log p_{nucl}\right\}\right) \quad (1)$$

#### B. (2) Computing P-value of best hit

All hits of each $q$ against the $N_t$ proteins in a specific phage genome $T$ are examined by their pairwise P-values, and the hit with the lowest pairwise P-value ("best hit") is retained. SpacePHARER computes the P-value of the best hit $p_{\text{bh}}(q)$ using first order statistics, i.e. the P-value of taking the minimum pairwise P-value ($p(q)$), given that a total of $N_t$ pairwise P-values were examined:

$$p_{\text{bh}}(q) = P\left(p\left(q\right) \leq p\right) = 1 - \left(1 - p\right)^{N_t} \quad (2)$$

#### C. (3) Combining P-values using a modified truncated product method

In this step, we aim to combine the evidence from several best hits between a spacer set $Q$ and a phage genome $T$. We sort the $p_{\text{bh}}$ of the given set $Q$ of $N_q$ sequences in ascending order and denote the $i$'th $p_{\text{bh}}$ as $p_i$. When combining independent P-values of individual hits, one needs to take into account the number of individual hits and the strength of each hit. The truncated product method combines independent P-values into a score by multiplying all $p_{\text{bh}}(q)$ smaller than a threshold $p_0$ [14],

$$S_{\text{comb}} = -\log \prod_{i=1}^{N_q} p_i^{I(p_i < p_0)}, \quad (3)$$

where $I(\cdot)$ is the indicator function that returns 1 if the argument is true and otherwise returns 0.

In SpacePHARER, we modified the truncated product method for better performance. We take the product of the smallest best-hit P-value $p_1$ times the ratio between $p_i$ and the threshold $p_0$ for all further $p_i$ below the threshold $p_0$:

$$S_{\text{comb}} = -\log\left(p_1 \times \prod_{i=2}^{N_q}\left(\frac{p_i}{p_0}\right)^{I(p_i<p_0)}\right) \quad (4)$$

For the threshold, we set $p_0 = 1/(N_q + 1)$, which corresponds to marginal significance, with an E-value of $N_q/(N_q + 1)$ just below 1. This ensures that the combined score for null model-distributed P-values $p_i$ only rarely gets boosted by a contribution from the second-best $p_i$.

#### D. (4) Determining true predictions

SpacePHARER predicts matches *de novo*, i.e. without relying on any known phage-host relationships, by controlling for estimated false discovery rate (FDR). The FDR is the proportion of false predictions among all predictions:

$$\text{FDR} = \frac{\text{FP}}{\text{FP} + \text{TP}} \quad (5)$$

We implemented an FDR estimation approach similar to that of the R package "fdrtool" [11]. In essence, we estimate the FDR by a Grenander decreasing density estimate of the empirical cumulative distribution function (ECDF). This non-parametric approach achieves its robustness by ensuring monotonicity of the FDR.

SpacePHARER uses a null model dataset to estimate the proportion of false predictions. The same search and statistical computation procedures described in Steps 1, 2 and 3 of the algorithm are performed on a given null model dataset, e.g. inverted phage ORFs or eukaryotic viral ORFs. Inverting target ORFs as null model dataset can be easily performed by specifying one parameter when preparing the input.

To compute an empirical P-value for each query spacer set $Q$, we sort for each $Q$ the combined scores $S_{\text{comb}}$ of matches in the original target dataset of phage proteomes in ascending order. For each $S_{\text{comb}}$ value in the target dataset, we calculate an empirical P-value $p_{\text{emp}}$ by using the fraction of $Q-T$ matches with a combined score that is below $S_{\text{comb}}$ in the null model dataset. We denote the number of $Q-T$ matches below the cutoff as $K$ and the total number of matches using the null model dataset as $N_{\text{null}}$. The empirical P-value is then computed as

$$p_{\text{emp}}(S_{\text{comb}}) = \frac{K + 0.5}{N_{\text{null}} + 1}, \quad (6)$$

where, to stabilize the estimate, we used half pseudocounts with P-values at 0 and 1. In the following, we abbreviate these empirical P-values as $p$, or $p_Q$ for query set $Q$.

If we knew the fraction $\pi_0$ of false positives among all $Q-T$ matches, we could in principle estimate the false discovery rate simply as

$$\text{FDR}(p) = \frac{\text{FP}_p}{(\text{TP} + \text{FP})_p} \approx \frac{p\,\pi_0}{F_{\text{emp}}(p)}, \quad (7)$$

where $p\,\pi_0$ is the fraction of false positives with empirical P-value less than $p_i$. $F_{\text{emp}}(p)$ is the empirical cumulative distribution function of the $p_Q$, in other words $F_{\text{emp}}(p)$ is the number of query sets $Q$ with best matches $p_Q \leq p$.

We can increase the robustness of the estimate by using the fact that the true probability distribution of P-values $f(p)$ must be monotonously decreasing. This will also ensure that the FDR decreases with increasing $p$, which is often violated with the simple procedure above. The Grenander estimate [11] is a simple, efficient procedure to obtain a robust estimate $\hat{F}(p)$ of $F(p)$ from $F_{\text{emp}}(p)$ that has monontonously decreasing density $\hat{f}(p) = d\hat{F}(p)/dp$. We simply obtain the convex hull of the area under the $F_{\text{emp}}(p)$ curve, that is, the smallest function $\hat{F}(p)$ with $\hat{F}(p) \geq F_{\text{emp}}(p)$ that yields a convex area under the curve. This results in a piecewise constant, monotonously decreasing density function $\hat{f}(p) = d\hat{F}(p)/dp$ with steps at points $p_i$ with $p_{\text{last}} = 1$. We estimate the proportion of true null hypotheses $\pi_0$ as the average density using the last two steps,

$$\pi_0 = \frac{\hat{F}(p_{\text{last}}) - \hat{F}(p_{\text{last}-2})}{p_{\text{last}} - p_{\text{last}-2}}. \quad (8)$$

Finally, we compute the estimated FDR corresponding to each empirical P-value $p$ (Fig.1A) as

$$\text{FDR}(p) = \frac{\text{FP}_p}{(\text{TP} + \text{FP})_p} = \frac{p\,\pi_0}{\hat{F}(p)}. \quad (9)$$

By default, SpacePHARER has an FDR cutoff of 0.05, and reports all matches in the test whose $S_{\text{comb}}$ corresponds to this FDR value or lower. Users can select other suitable FDR cutoffs to retain more or fewer predictions.

#### E. (5) Scanning for possible PAMs

For some CRISPR-Cas systems, protospacer adjacent motifs (PAMs) are required for the recognition of foreign invader sequences. After reporting phage-host pairs and their hits, SpacePHARER can perform a scan for possible PAMs. For this, SpacePHARER by default extracts 10 nt long fragments flanking the matched protospacer region at the 5' and 3' side, in guide-centric orientation (PAM is located on the strand that matched the spacer sequence). Users can increase or decrease the length of the flanking sequence. Both the 5' and 3' flanking sequences are searched in a list of consensus PAM patterns from representative CRISPR-Cas systems [5]. Since many CRISPR detection tools cannot reliably predict the orientation of the CRISPR array, the 5' and 3' flanking sequences on the reverse strand are also searched and two additional possible PAMs are reported. Users should refer to all possible PAMs without the accurate orientation information of the array.

### II. OPTIMIZING PARAMETERS FOR SHORT FRAGMENTS SEARCH

Different substitution matrices are optimal for comparing sequences that have diverged to different degrees. By default, MMseqs2 search [10] uses the BLOSUM62 matrix with standard gap penalties: gap open cost of 11 and gap extend cost of 1 , which is more suited for long alignments and detecting weak protein similarities. Conversely for shorter sequences and higher protein similarity, one should consider a "shallower" (higher bit score per aligned column) matrix, and higher gap penalties to prevent gaps [9]. Searching with VTML40 matrix [8] with gap open cost of 16 and gap extend cost of 2 yielded the highest sensitivity with 20% our test dataset at FDR cutoff of 0.05 (Figure S2). We introduced a series of VTML matrices in MMseqs2 to solve general problems of short sequence search. After introducing the additional nucleotide alignment step, the search parameter combination (VTML40 matrix, gap open cost of 16 and gap extend cost of 2) remains the highest in sensitivity (result not shown).

### III. PREDICTING MATCHES USING BLASTN

We compared SpacePHARER's performance with the state-of-the-art method using BLASTN. To generate a comparable result, we performed the search step with BLASTN and the downstream FDR control with SpacePHARER. We used BLASTN [1] to first query the 80% test spacer dataset against 7,824 phage genomes, then against 7,824 inverted phage genomes or 11,304 eukaryotic viral genomes as a null model database. For all searches we used the parameters: -max_target_seqs 10000000 -dust no -word_size 7 -outfmt '6 std qcovs' and recorded the running time. Hits with at least 95% sequence identity and 95% query coverage (i.e., one or two mismatches were allowed) were retained. We grouped the hits into matches (unique phage-host genome pairs) and retained the minimum pairwise E-value of the hits. We sorted the pairwise E-values of hits in ascending order for both searches and counted the matches at a given pairwise E-value cutoff. Therefore, we could calculate an FDR in the same way SpacePHARER does (described in section I.D) and compare the number of true predictions produced by the two methods (Figure 1B).

At FDR = 0.05, SpacePHARER predicted 2 and 1.5× more matches than BLASTN using 90% and 85% sequence identity and query coverage cutoffs (i.e allowing up to 4 and 6 mismatches, respectively) (Figure S2).

### IV. HOST TAXONOMIC RANK ANALYSIS

To assess the sensitivity of SpacePHARER at different host taxonomic rank, we searched with CRISPR spacers extracted from 1,066 bacterial genomes against 809 phage genomes with annotated host taxonomy [4], then against inverted ORFs of the 809 phage genomes as null model dataset. For each phage, SpacePHARER predicted the host lowest common ancestor (LCA) based on an weighted LCA procedure [7].

We demanded a stricter FDR cutoff of 0.02 for matches that should be taken into account for the host taxonomic rank prediction. In order to limit the number of false taxonomic prediction due to incomplete databases, the LCA result was further corrected according to the average nucleotide sequence identity of the reported matches [6]. We used the following cutoffs for maximal taxonomic resolution: > 86% (species), > 84% (genus), > 82% (family), > 80% (order), > 78% (class), > 76% (phylum), > 74% (kingdom). Lower values were assigned at the superkingdom level. The taxonomic FDR cutoff and sequence identity cutoffs are user-definable parameters for the weighted LCA procedure.

We performed the same search using BLASTN with parameters: blastn-short -dust no -word_size 7 -outfmt '6 std qcovs' -evalue 1 -gapopen 10 -gapextend 2 -penalty -1 [4]. Hits with at least 95% sequence identity and 95% query coverage were retained. For each phage, the bacterium with the lowest pairwise E-value was predicted to be its host.

For ranks lower than phylum, we only included the predictions with the taxonomic resolution of the respective rank or below. At the species level, SpacePHARER predicted 142/237 hosts (60%), comparing with 112/232 hosts of BLASTN (48%). SpacePHARER predicted the correct host for more phages at all taxonomic ranks, while including most of the BLASTN predictions on the same rank and sometimes even those agreeing only on a higher rank(Figure 1C, Figure S3).

### V. IDENTIFYING MIS-ANNOTATIONS IN EUKARYOTIC VIRAL DATASET

Throughout this study we used the set of *eukaryotic viral genomes* as a null model dataset, assuming any match between a prokaryotic genome and a eukaryotic virus is false. Here, we used SpacePHARER's second mode of FDR control to detect viruses that were potentially misannotated as eukaryotic viruses. To that end, we first ran the SpacePHARER workflow with the full spacer dataset against the *eukaryotic viral genomes* as the target database, and then, against inverted *eukaryotic viral* ORFs as the null model database. We used the null set to estimate the FDR as described in section I.D.

By applying the same FDR cutoff of 0.05, we identified 11 viruses out of the 11,304 that matched a prokaryotic host (yielding a total of 12 matches). We observed three groups within these matches. The first group consisted of two matches between the smacovirus family (KP264966.1 and KY086299.1) and the archaeon CP005934.1 (*Candidatus Methanomassiliicoccus intestinalis*). Indeed this family has been recently reported as mis-annotated as "eukaryotic virus" by Díez-Villaseñor and Rodriguez-Valera [3]. The second group consisted of two matches between KT809302.1 (Haloarcula californiae icosahedral virus 1) and family *Halobacteriaceae* (CP001687.1 and LIST01000008.1). These matches are likely due to mis-annotation of the virus as "eukaryotic virus". The labeled host of this virus is *Haloarcula californiae*, which is an archaeon that belongs to the same family as our matches. The third group consisted of 8 members of the genus *Mimivirus* that were matched to HE978663.1 (*Ruminococcus sp.* JC304) and JAAF01000022.1 (*Fusobacterium necrophorum* DAB). Table I shows the standard output from SpacePHARER of this search. We suspect the matches of the third group are due to spacer mis-annotation and do not represent a real virus-host relationship. It was previously reported that Mimiviruses acquire bacterial genes, even of the class *Clostridia* [12][13]. In the case of *Ruminococcus sp.* JC304, when we inspected the bacterial genomic region from which the spacers were extracted, we found that the entire region is likely to be a full bacterial ORF, rather than a CRISPR array. Thus, we conclude that in these cases, the mis-annotation is of the CRISPR array, rather than of the virus.

### VI. SOFTWARE VERSIONS

| Name | Version |
|---|---|
| SpacePHARER | Git: 1d1f1b2 |
| BLASTN | 2.9.0+ |

TABLE II. Software versions used in this manuscript.


[1] Altschul, S.F. et al (1990). Basic local alignment search tool. *J. Mol. Biol.*, **215**(3), 403–410.

[2] Brunson, J.C. (2020). ggalluvial: Layered grammar for alluvial plots. *J. Open Source Softw.*, **5**(49), 2017.

[3] Díez-Villaseñor, C. and Rodriguez-Valera, F. (2019). CRISPR analysis suggests that small circular single-stranded dna smacoviruses infect archaea instead of humans. *Nat. Commun.*, **10**(1), 294.

[4] Edwards, R.A. et al (2015). Computational approaches to predict bacteriophage–host relationships. *FEMS Microbiol. Rev.*, **40**(2), 258–272.

[5] Leenay, R.T. and Beisel, C.L. (2017). Deciphering, communicating, and engineering the crispr pam. *Journal of molecular biology*, **429**(2), 177–191.

[6] Levy Karin, E. et al (2020). Metaeuk—sensitive, high-throughput gene discovery, and annotation for large-scale eukaryotic metagenomics. *Microbiome*, **8**(1), 48.

[7] Mirdita, M. et al (2020). Fast and sensitive taxonomic assignment to metagenomic contigs. *bioRxiv.* doi:10.1101/2020.11.27.401018.

[8] Müller, T. et al (2002). Estimating amino acid substitution models: A comparison of Dayhoff's estimator, the resolvent approach and a maximum likelihood method. *Mol. Biol. Evol.*, **19**(1), 8–13.

[9] Pearson, W.R. (2013). Selecting the Right Similarity-Scoring Matrix. *Current Protocols in Bioinformatics*, **43**(1), 3.5.1–3.5.9.

[10] Steinegger, M. and Söding, J. (2017). MMseqs2 enables sensitive protein sequence searching for the analysis of massive data sets. *Nat. Biotechnol.*, **35**(11), 1026–1028.

[11] Strimmer, K. (2008). A unified approach to false discovery rate estimation. *BMC Bioinformatics*, **9**(1), 303.

[12] Yoshida, T. et al (2011). Mimivirus reveals mre11/rad50 fusion proteins with a sporadic distribution in eukaryotes, bacteria, viruses and plasmids. *Virology journal*, **8**, 427–427.

[13] Yutin, N. et al (2014). Origin of giant viruses from smaller dna viruses not from a fourth domain of cellular life. *Virology*, **466-467**, 38 – 52. Special issue: Giant Viruses.

[14] Zaykin, D. et al (2002). Truncated product method for combining p-values. *Genet. Epidemiol.*, **22**(2), 170–185.

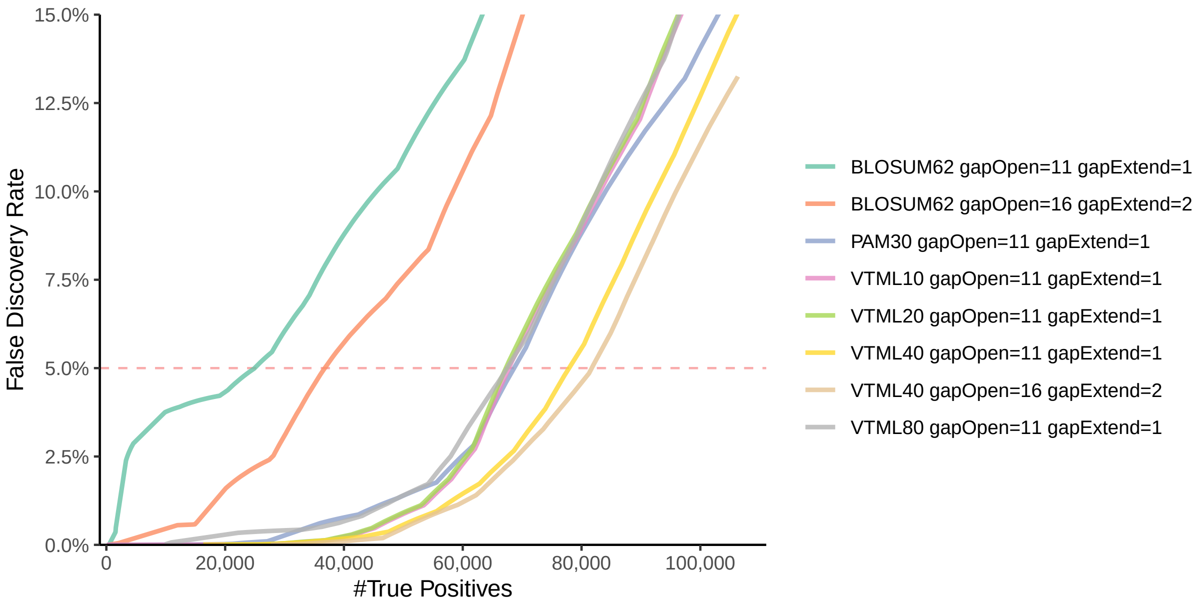


FIG. 1. Performance comparison of SpacePHARER with different search parameters (substitution matrix and gap penalties), evaluated by the number of true positive (TP) predictions at different false discovery rates (FDRs). Predictions were made by using a optimization spacer dataset (6,067 genomes, 20% of all prokaryotic genomes) against a database of 7,824 phage genomes, with inverted phage ORFs as null model database. Searching with VTML40 matrix with gap open (16) and gap extend (2), among various combinations of substitution matrix and gap penalties, yields most true positive matches than any other parameter combination at FDR cutoff of 0.05.

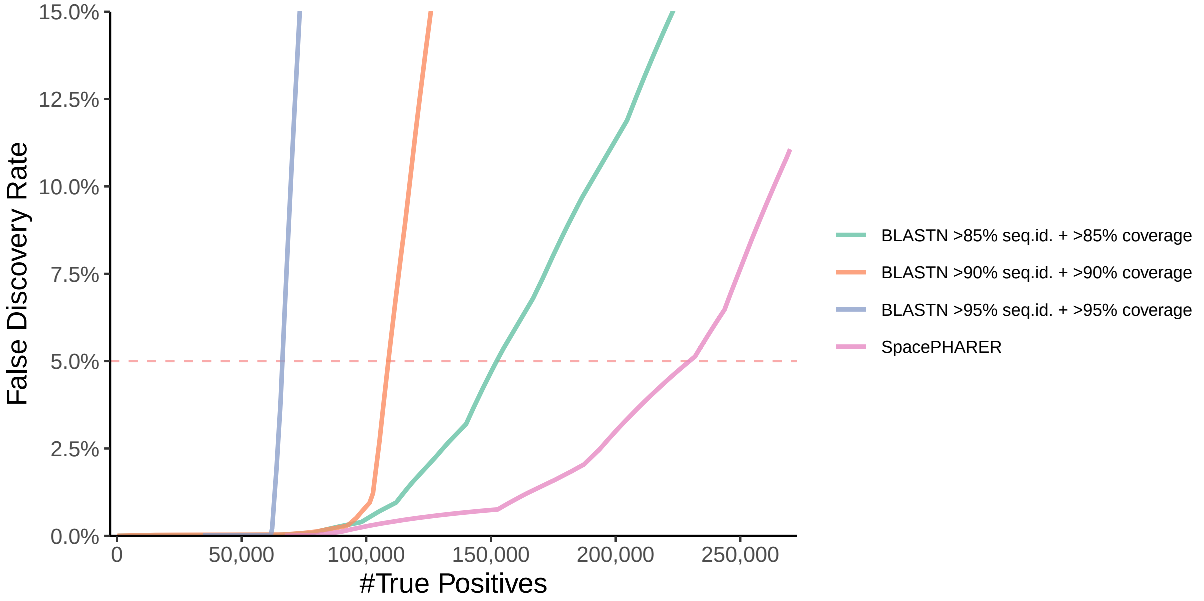


FIG. 2. Performance comparison of SpacePHARER with BLASTN using different sequence identity and query coverage cutoffs (95%, 90% and 85%), evaluated by the number of true positive (TP) predictions at different false discovery rates (FDRs). Predictions were made by using a spacer test dataset (24,322 genomes, 80% of all prokaryotic genomes) against a database of 7,824 phage genomes, with inverted phage ORFs as null model database. (Note that the FDR control procedure developed for SpacePHARER is not standard for BLASTN and has been applied here only for the purpose of FDR analysis.)

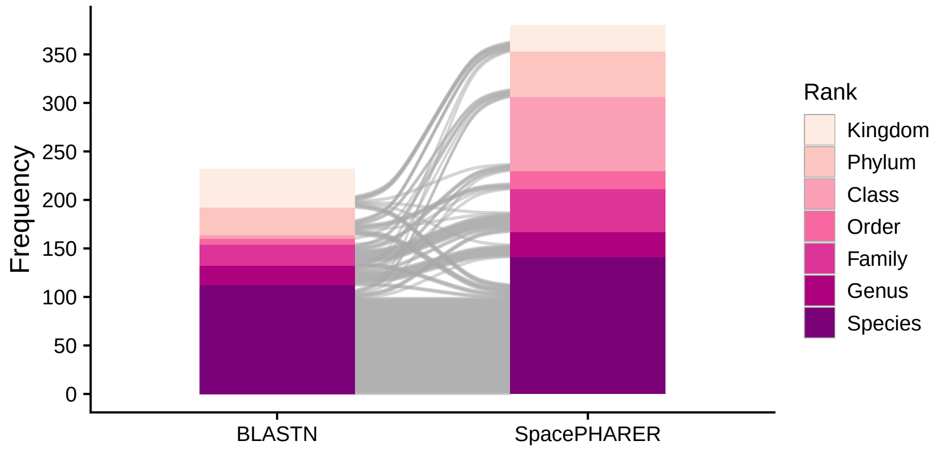


FIG. 3. Performance comparison of BLASTN (left) and SpacePHARER (right), evaluated by the number of host predictions that agree with annotated host taxonomy at different taxonomic ranks. The grey alluvia [2] represent the host predictions that were made by both SpacePHARER and BLASTN. Predictions were made using a validation spacer dataset (1,066 genomes) against a validation database of 809 phage genomes with annotated host taxonomy. SpacePHARER prediction was further corrected with inverted phage ORFs as null model database, and FDR cutoff of 0.02.

<table>
<tr><td>#CP005934.fas</td><td>KP264966.1</td><td>7.588E+01</td><td>6</td><td></td><td></td><td></td><td></td><td></td></tr>
<tr><td>>CP005934.1_930280_937725_19_spacer_931524_35</td><td>KP264966.1</td><td>1.023E-04</td><td>35</td><td>3</td><td>546</td><td>578</td><td>CCT|-</td><td>-|AGG</td></tr>
<tr><td>>CP005934.1_930280_937725_20_spacer_931590_37</td><td>KP264966.1</td><td>2.833E-04</td><td>1</td><td>36</td><td>2241</td><td>2206</td><td>CCT|-</td><td>-|AGG</td></tr>
<tr><td>>CP005934.1_930280_937725_23_spacer_931792_37</td><td>KP264966.1</td><td>1.034E-09</td><td>1</td><td>36</td><td>1821</td><td>1786</td><td>CCA|-</td><td>-|TGG</td></tr>
<tr><td>>CP005934.1_930280_937725_24_spacer_931860_37</td><td>KP264966.1</td><td>3.121E-07</td><td>3</td><td>35</td><td>606</td><td>638</td><td>CCA|TGG</td><td>-|TGG</td></tr>
<tr><td>>CP005934.1_930280_937725_25_spacer_931928_36</td><td>KP264966.1</td><td>3.399E-13</td><td>1</td><td>36</td><td>2161</td><td>2126</td><td>CCG|-</td><td>-|CGG</td></tr>
<tr><td>>CP005934.1_930280_937725_25_spacer_931928_36</td><td>KP264966.1</td><td>1.713E-11</td><td>2</td><td>34</td><td>2160</td><td>2128</td><td>CCG|-</td><td>-|CGG</td></tr>
<tr><td>#CP005934.fas</td><td>KY086299.1</td><td>5.640E+01</td><td>4</td><td></td><td></td><td></td><td></td><td></td></tr>
<tr><td>>CP005934.1_930280_937725_19_spacer_931524_35</td><td>KY086299.1</td><td>6.205E-04</td><td>35</td><td>3</td><td>1922</td><td>1890</td><td>CCT|-</td><td>-|AGG</td></tr>
<tr><td>>CP005934.1_930280_937725_23_spacer_931792_37</td><td>KY086299.1</td><td>3.399E-13</td><td>2</td><td>37</td><td>641</td><td>676</td><td>CCA|-</td><td>-|AGG</td></tr>
<tr><td>>CP005934.1_930280_937725_20_spacer_931590_37</td><td>KY086299.1</td><td>4.613E-05</td><td>1</td><td>36</td><td>220</td><td>255</td><td>CCT|-</td><td>-|TGG</td></tr>
<tr><td>>CP005934.1_930280_937725_23_spacer_931792_37</td><td>KY086299.1</td><td>3.399E-13</td><td>1</td><td>36</td><td>640</td><td>675</td><td>CCA|-</td><td>-|TGG</td></tr>
<tr><td>#LIST01000008.fas</td><td>KT809302.1</td><td>1.295E+01</td><td>1</td><td></td><td></td><td></td><td></td><td></td></tr>
<tr><td>>LIST01000008.1_120573_126312_45_spacer_123484_36</td><td>KT809302.1</td><td>2.376E-06</td><td>36</td><td>1</td><td>22375</td><td>22340</td><td>-|-</td><td>TTC|-</td></tr>
<tr><td>#CP001687.fas</td><td>KT809302.1</td><td>2.639E+01</td><td>2</td><td></td><td></td><td></td><td></td><td></td></tr>
<tr><td>>CP001687.1_1415738_1419119_25_spacer_1417344_34</td><td>KT809302.1</td><td>1.137E-07</td><td>6</td><td>32</td><td>6826</td><td>6852</td><td>-|CAAGAA</td><td>-|ACGGGATT</td></tr>
<tr><td>>CP001687.1_1415738_1419119_25_spacer_1417344_34</td><td>KT809302.1</td><td>1.137E-07</td><td>32</td><td>6</td><td>6852</td><td>6826</td><td>-|CAAGAA</td><td>-|ACGGGATT</td></tr>
<tr><td>#HE978663.fas</td><td>JN258408.1</td><td>1.054E+02</td><td>2</td><td></td><td></td><td></td><td></td><td></td></tr>
<tr><td>>HE978663.1_7481_7851_2_spacer_7588_70</td><td>JN258408.1</td><td>1.755E-28</td><td>2</td><td>70</td><td>806538</td><td>806606</td><td>TTC|-</td><td>-|-</td></tr>
<tr><td>>HE978663.1_7481_7851_4_spacer_7765_58</td><td>JN258408.1</td><td>5.707E-20</td><td>2</td><td>58</td><td>806715</td><td>806771</td><td>TTC|-</td><td>-|-</td></tr>
<tr><td>#HE978663.fas</td><td>JX885207.1</td><td>9.775E+01</td><td>2</td><td></td><td></td><td></td><td></td><td></td></tr>
<tr><td>>HE978663.1_7481_7851_2_spacer_7588_70</td><td>JX885207.1</td><td>1.755E-28</td><td>2</td><td>70</td><td>767273</td><td>767341</td><td>TTC|-</td><td>-|-</td></tr>
<tr><td>>HE978663.1_7481_7851_4_spacer_7765_58</td><td>JX885207.1</td><td>1.187E-16</td><td>2</td><td>58</td><td>767450</td><td>767506</td><td>TTC|-</td><td>-|-</td></tr>
<tr><td>#HE978663.fas</td><td>KF527229.1</td><td>8.186E+01</td><td>2</td><td></td><td></td><td></td><td></td><td></td></tr>
<tr><td>>HE978663.1_7481_7851_1_spacer_7510_49</td><td>KF527229.1</td><td>2.905E-18</td><td>2</td><td>49</td><td>935992</td><td>935945</td><td>TTC|-</td><td>-|-</td></tr>
<tr><td>>HE978663.1_7481_7851_4_spacer_7765_58</td><td>KF527229.1</td><td>5.707E-20</td><td>2</td><td>58</td><td>935836</td><td>935780</td><td>TTC|-</td><td>-|-</td></tr>
<tr><td>#HE978663.fas</td><td>KU877344.1</td><td>9.775E+01</td><td>2</td><td></td><td></td><td></td><td></td><td></td></tr>
<tr><td>>HE978663.1_7481_7851_2_spacer_7588_70</td><td>KU877344.1</td><td>1.755E-28</td><td>2</td><td>70</td><td>780352</td><td>780420</td><td>TTC|-</td><td>-|-</td></tr>
<tr><td>>HE978663.1_7481_7851_4_spacer_7765_58</td><td>KU877344.1</td><td>1.187E-16</td><td>2</td><td>58</td><td>780529</td><td>780585</td><td>TTC|-</td><td>-|-</td></tr>
<tr><td>#HE978663.fas</td><td>JX975216.1</td><td>1.015E+02</td><td>2</td><td></td><td></td><td></td><td></td><td></td></tr>
<tr><td>>HE978663.1_7481_7851_2_spacer_7588_70</td><td>JX975216.1</td><td>1.755E-28</td><td>2</td><td>70</td><td>781866</td><td>781934</td><td>TTC|-</td><td>-|-</td></tr>
<tr><td>>HE978663.1_7481_7851_4_spacer_7765_58</td><td>JX975216.1</td><td>2.682E-18</td><td>2</td><td>58</td><td>782043</td><td>782099</td><td>TTC|-</td><td>-|-</td></tr>
<tr><td>#HE978663.fas</td><td>MG779360.1</td><td>1.093E+02</td><td>2</td><td></td><td></td><td></td><td></td><td></td></tr>
<tr><td>>HE978663.1_7481_7851_2_spacer_7588_70</td><td>MG779360.1</td><td>3.447E-30</td><td>2</td><td>70</td><td>9786</td><td>9854</td><td>-|-</td><td>-|-</td></tr>
<tr><td>>HE978663.1_7481_7851_4_spacer_7765_58</td><td>MG779360.1</td><td>5.707E-20</td><td>2</td><td>58</td><td>9963</td><td>10019</td><td>-|-</td><td>-|-</td></tr>
<tr><td>#HE978663.fas</td><td>JN885991.1</td><td>4.536E+01</td><td>2</td><td></td><td></td><td></td><td></td><td></td></tr>
<tr><td>>HE978663.1_7481_7851_3_spacer_7687_49</td><td>JN885991.1</td><td>2.061E-02</td><td>2</td><td>46</td><td>497977</td><td>498021</td><td>CCT|-</td><td>TTG|AGG</td></tr>
<tr><td>>HE978663.1_7481_7851_4_spacer_7765_58</td><td>JN885991.1</td><td>5.707E-20</td><td>2</td><td>58</td><td>498055</td><td>498111</td><td>-|-</td><td>-|-</td></tr>
<tr><td>#JAAF01000022.fas</td><td>KY684109.1</td><td>1.295E+01</td><td>1</td><td></td><td></td><td></td><td></td><td></td></tr>
<tr><td>>JAAF01000022.1_41_3914_26_spacer_1726_36</td><td>KY684109.1</td><td>2.376E-06</td><td>1</td><td>36</td><td>185359</td><td>185324</td><td>TCT|TGAAGTTT</td><td>TCA|-</td></tr>
</table>

TABLE I. Sample output format of SpacePHARER, demonstrated by matches when searching the full spacer dataset against eukaryotic viral ORFs as a target database and inverted eukaryotic viral ORFs as null model database. Each match line starts with "#", followed by the prokaryote accession (the file from which spacers were extracted), viral genome accession, $S_{comb}$ and the number of hits in the match. Each hit line starts with ">", followed by the spacer sequence header, viral genome accession, $p_{bh}$, spacer start, spacer end, viral genome start, viral genome end, and the possible PAM sequences on forward and reverse strand (5'|3'). Additionally (not shown), the aligned sequences can be printed following each hit line
